# A low-cost and high-precision scanning electrochemical microscope built with open source tools

**DOI:** 10.1101/645283

**Authors:** Alperen Guver, Nafetalai Fifita, Peker Milas, Michael Straker, Michael Guy, Kara Green, Taha Yildirim, Ilyas Unlu, Veysel Yigit, Birol Ozturk

## Abstract

A low-cost Scanning Electrochemical Microscope (SECM) was built with a 0.6 pA current measurement capability potentiostat and submicron resolution motorized stage, using open source software and hardware tools. The high performance potentiostat with a Python graphical user interface was built based on an open source project. Arduino boards, stepper motors, a manual XY micromanipulator stage, 3D printed couplers and gears were used in building the motorized stage. An open source motor control software was used for moving the motorized stage with high precision. An inverted microscope was utilized for viewing a standard microelectrode while scanning. The setup was tested in the formation of a map of electrochemical signals from an array of pores on a parafilm membrane. As the setup will be used in future biosensing experiments, DNA hybridization detection experiments were also performed with the setup.

## Introduction

There is an increasing interest in manufacturing custom laboratory research instruments with the simplified tools developed by the open source community. This approach has been spurred due to high cost and resulting lack of accessibility to high performance laboratory equipment in certain education and research institutions. The reduced cost and increasing availability of 3D printers and easy to program electronic boards are playing a key role in motivating researchers to build their own lab instruments [1]. Chagas and co-workers remarkably developed a whole open-source 3D printable platform for fluorescence microscopy, optogenetics and accurate temperature control which costs 100 Euros to build [2]. Various groups have been successful in the development of field compatible inexpensive potentiostats which work with smartphone applications [3,4]. Meloni and co-workers developed a 3D printed scanning electrochemical microscope (SECM) for a total cost of one hundred dollars [5]. A 5-micron resolution motorized stage was built from 3D printed parts that was employed in a screening microscopy [6]. Furthermore, this DIY approach not only provides an innovative solution to the lack of instrumentation accessibility issue, but also is training opportunity for students to gain and develop design and troubleshooting skills during the process of building instruments [1].

Scanning Electrochemical Microscopy (SECM) is a powerful analytical tool for the identification of local electrochemical processes at various interfaces between gases, liquids and solids [7-10]. The commercially available SECMs are capable of carrying out nanometer resolution scans and sub picoampere current measurements. However, they cost several tens of thousand dollars. Here, we describe the procedure for building a high performance SECM with an inverted optical microscope, a custom-built motorized stage and a DSTAT potentiostat with 0.6 pA current measurement capability, where the cost of the custom-built parts was less than $250. Fig 1 shows a picture of the custom-built SECM with individual elements labeled. The setup has been successfully tested in standard electrochemical measurements, in the formation of an electrochemical signal image through scanning an array of pores on a membrane and in DNA hybridization experiments.

**Fig 1.**
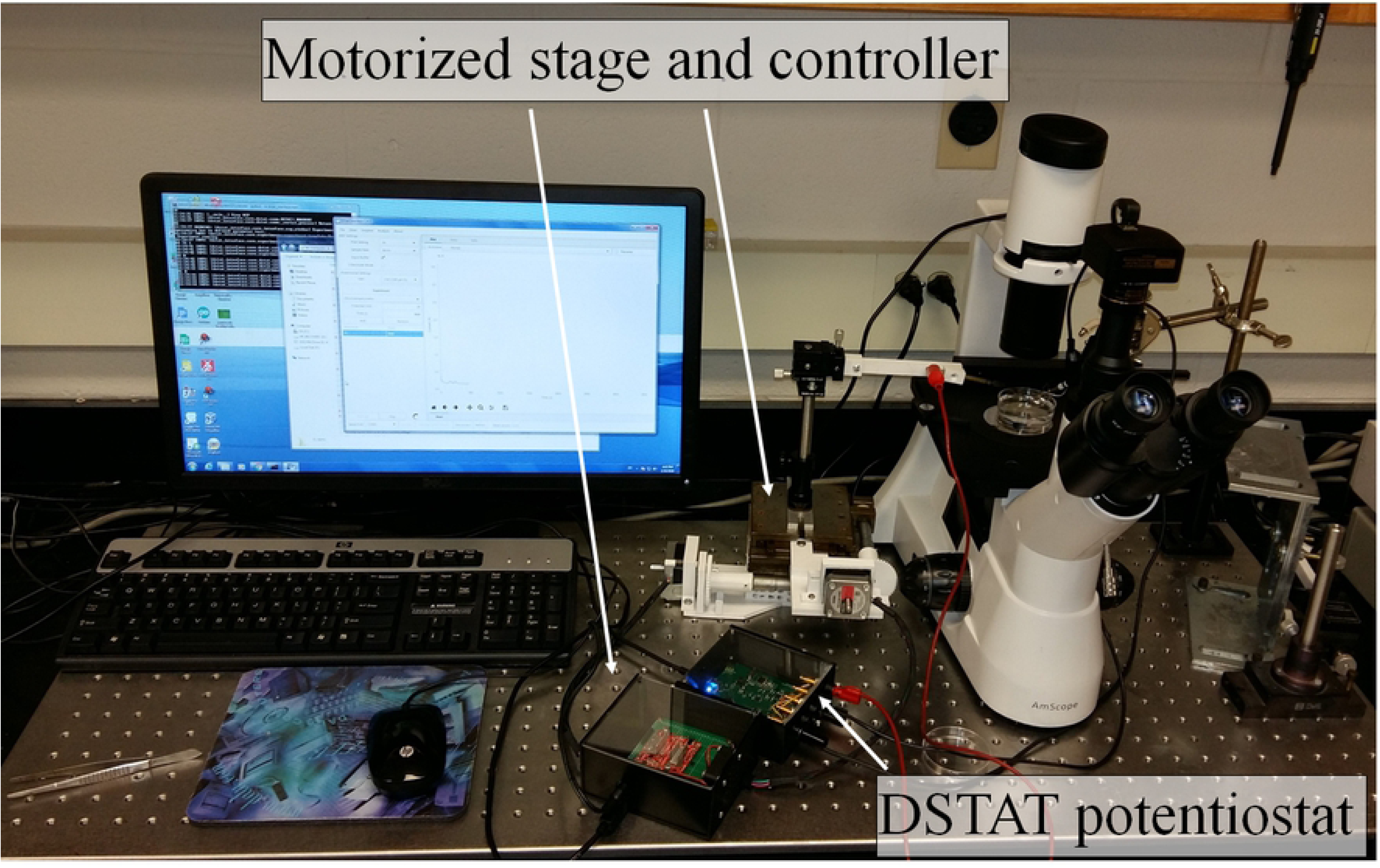
SECM setup image. An image of the custom built SECM setup with custom built elements labeled.

## Potentiostat Selection

A potentiostat is the core instrument of an SECM and its performance is a limiting factor for the type of measurements that can be conducted with the SECM. For example, battery research requires high voltage output but not low current measurements. The custom built SECM in this project will be used in the electrochemical detection of biomolecules, thus the low-level current detection capability was the determining factor in the selection of the potentiostat. Moreover, the overarching goal of the project was to custom build a low-cost SECM. However, the price range for commercially available standard potentiostats with low-current measurement capability is $2,000 - $20,000. There have been several attempts to significantly reduce the cost of the potentiostat with DIY approach, using open source programmable Arduino boards [11,12] and with other custom circuit board designs [4,4,13-15]. As shown in Table 1, the cost of Arduino based potentiostats [11,12] are in the $30-40 range but they can only measure high microampere currents and they don’t have the square wave voltage (SWV) measurement capability, which is required for the detection of low concentration analytes. Custom design circuit board based UWED and uMED potentiostats [4,4] have the advantage of having small-form factors as they are built for field applications, where cell phones and apps are used as interfaces. These potentiostats can also only measure high microampere currents. Other custom built potentiostats offer low microampere measurement capability and their cost range is $80-100 [14,14]. In building the custom SECM in this project, the DSTAT potentiostat was chosen due to its superior low current (600fA) measurement capability [15] and it still has a moderate cost of around $120.

**Table 1.**
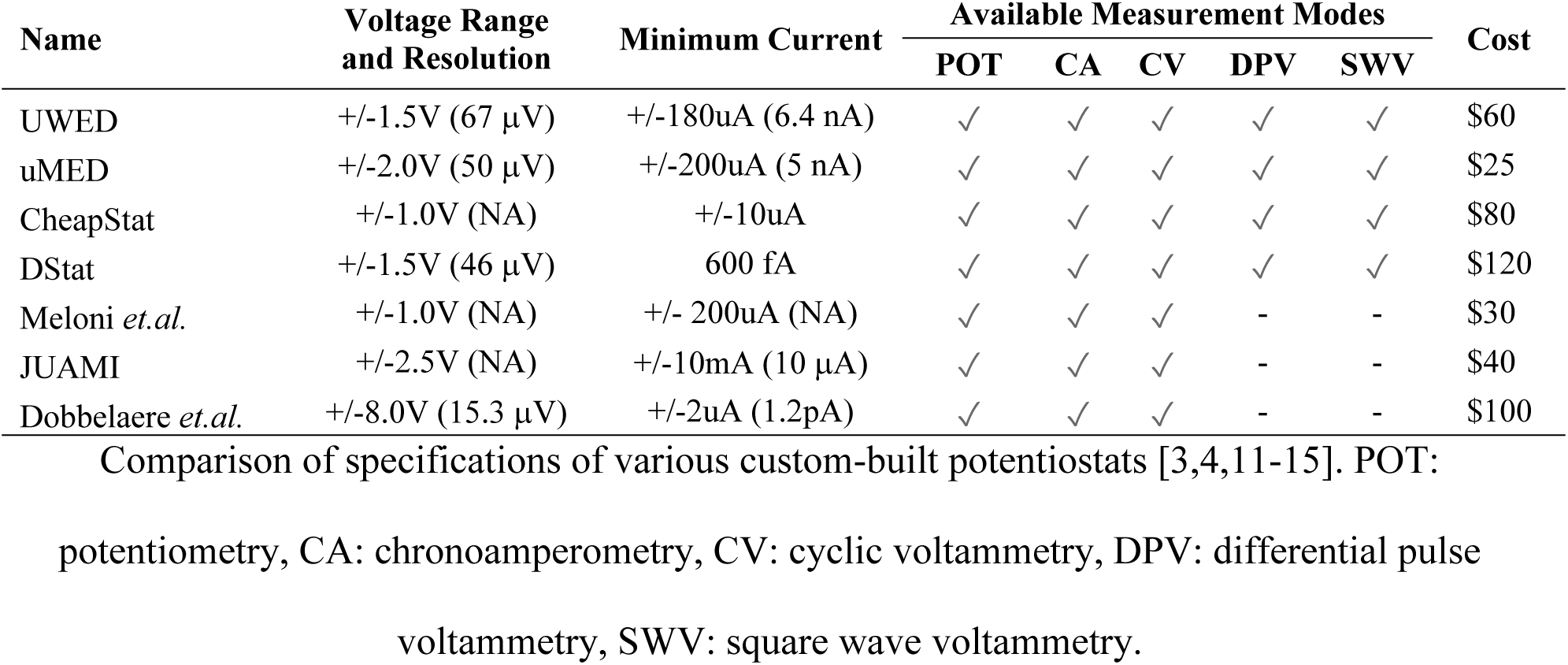
Specifications of different potentiostats.

| Name | Voltage Range and Resolution | Minimum Current | Available Measurement Modes |  |  |  |  | Cost |
| --- | --- | --- | --- | --- | --- | --- | --- | --- |
|  |  |  | POT | CA | CV | DPV | SWV |  |
| UWED | +/-1.5V (67 μV) | +/-180uA (6.4 nA) | ✓ | ✓ | ✓ | ✓ | ✓ | \$60 |
| uMED | +/-2.0V (50 μV) | +/-200uA (5 nA) | ✓ | ✓ | ✓ | ✓ | ✓ | \$25 |
| CheapStat | +/-1.0V (NA) | +/-10uA | ✓ | ✓ | ✓ | ✓ | ✓ | \$80 |
| DStat | +/-1.5V (46 μV) | 600 fA | ✓ | ✓ | ✓ | ✓ | ✓ | \$120 |
| Meloni *et.al.* | +/-1.0V (NA) | +/- 200uA (NA) | ✓ | ✓ | ✓ | - | - | \$30 |
| JUAMI | +/-2.5V (NA) | +/-10mA (10 μA) | ✓ | ✓ | ✓ | - | - | \$40 |
| Dobbelaere *et.al.* | +/-8.0V (15.3 μV) | +/-2uA (1.2pA) | ✓ | ✓ | ✓ | - | - | \$100 |

Comparison of specifications of various custom-built potentiostats [3,4,11-15]. POT:
potentiometry, CA: chronoamperometry, CV: cyclic voltammetry, DPV: differential pulse voltammetry, SWV: square wave voltammetry.

The DSTAT potentiostat was built by following the detailed instructions provided by the developers in their publication, supplementary materials and the online project website. Minor modifications were done to the 3D printed box and the new design is provided as a supplementary material to this manuscript. The cost of building the DSTAT potentiostat was around $160, which is similar to the originally provided estimate by the developers. As shown in Table 1, DSTAT potentiostat is capable of measuring sub picoampere signals, provides many different measurement modes, and its Python based user interface is very easy to use. An undergraduate student was able to build it by following clear directions in its publication [15] and its project website. Picoampere level currents were consistently measured with the custom built DSTAT in various low-level signal experiments and some of them will be presented below.

## Motorized Stage

The reproducible scanning property of an SECM depends on its high precision motorized stage. Commercially available SECMs have nanoscale scan step capability through the use piezo motor stages, however, commercial piezo motor stages have starting prices of several thousand dollars and there is no established DIY approach literature on building piezo motors. Stepper motor-controlled stages provide micron scale resolution at a much lower cost, which is sufficient for the goals of this project. We have built a stepper motor-controlled stage with submicron step size and 2.5cm range in both directions, using a manual XY stage, an Arduino board, two stepper motor driver shields, 3D printed parts and an open source user interface. An existing micrometer controlled manual XY stage was utilized in building the motorized stage, which can be purchased for $100 from various vendors. Others also reported on DIY approaches to building micromanipulators [16]. Fig 2 shows a close-up image of the custom-built motorized XY stage.

**Fig 2.**
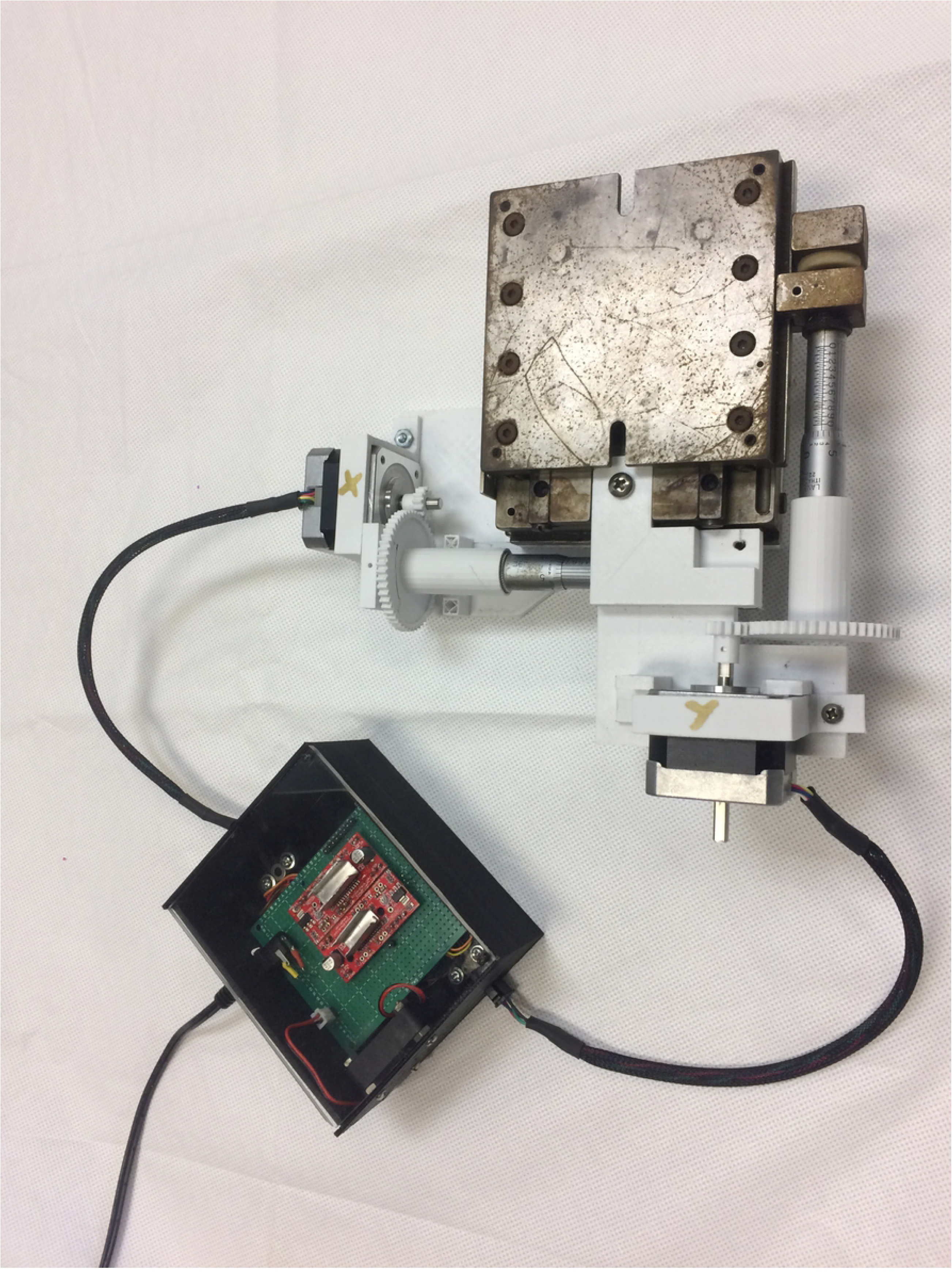
A close-up image of the motorized stage. The custom-built motorized stage parts including the manual XY stage, controller box, stepper motors, and 3D printed gears are shown.

As shown in the figure, the two-stepper motors were coupled to the manual XY stage via 3D printed parts and gears, where the difference in size and teeth numbers between gears enabled reduction of motor speed and hence the step size of the stage motion. The gear ratio was *N*_*Large-50*_/*N*_*Small-13*_ = 3.84, providing about 4 times reduction on the angular speed of each axis according to *w*_*L*_*N*_*L*_=*w*_*s*_*N*_*s*_. The stepper motors were controlled with the Arduino board in conjunction with Easy Driver v4.4 shields, which resulted in further reduction of motor speeds by enabling adjustments to the supplied currents to the stepper motors. The reduction of the stepper motor speed through gears and current control enabled stepwise motion of an axis by 500 nm in each step. This was demonstrated in the S1_Video as a supplementary information, where the tip of a tapered tungsten wire, that is attached to the motorized stage, covers the 10 micron distance between two lines on a calibration slide in 20 steps.

The stepper motor controller electronics including the Arduino board was housed in a custom 3D printed box. A custom Arduino shield was built using a perfboard to hold the two Easy Driver shields, the output power jacks for stepper motors and the DC power input jack, where a 9.75V DC adapter was utilized to power stepper drivers. A 5V fan was also installed in the box to cool down the Easy Driver shields during operation. The open source GRBL software with a graphical user interface was utilized in sending commands to the stage for stepwise motorized scan of a preferred area [17]. This software is also capable of automatically moving the XY stage according to a user uploaded image pattern and an example is presented as a supplementary information movie (S2_Video). GRBL software’s website provides detailed instructions for establishing communication between the software and the Arduino board [17]. A detailed parts list, 3D printable part files, GRBL software operation instructions, and a summary cost break down for the motorized stage are provided as supplementary information. We have also demonstrated controlling the motorized stage Arduino board through Python commands and in future upgrades, the motorized stage commands will be sent to the Arduino board motor controller through a single Python based interface, which will control both the motorized stage and the DSTAT potentiostat.

Once built, DSTAT potentiostat and the motorized stage can be used with any standard optical microscope to perform SECM experiments. An inverted microscope with enough clearance on top of its stage is preferred to accommodate the electrodes. The motorized stage is a modular bench top instrument and controls the motion of the working electrode of the DSTAT with extended arms to the top of the microscope stage as shown in Fig 1.

## SECM testing electrochemical signal mapping

The SECM was tested in forming an electrochemical signal image by scanning an array of 16 pores on a parafilm membrane as shown in Fig 3a. A membrane was formed by stretching parafilm over a 2.5cm × 2.5cm × 5mm container filled with 5mM K_3_Fe(CN)_6_ solution. The array of pores was formed by piercing the parafilm membrane with an electrochemically etched tungsten wire, which was prepared according to a previously established method in our lab [18]. Briefly, a 250 micron diameter tungsten wire (A-M Systems) was immersed in 2M NaOH solution in an oscillating fashion at 2 Hz frequency while applying 14V DC potential between the tungsten wire and a counter electrode that resides in the NaOH solution until a tapered tip is formed.

**Fig 3.**
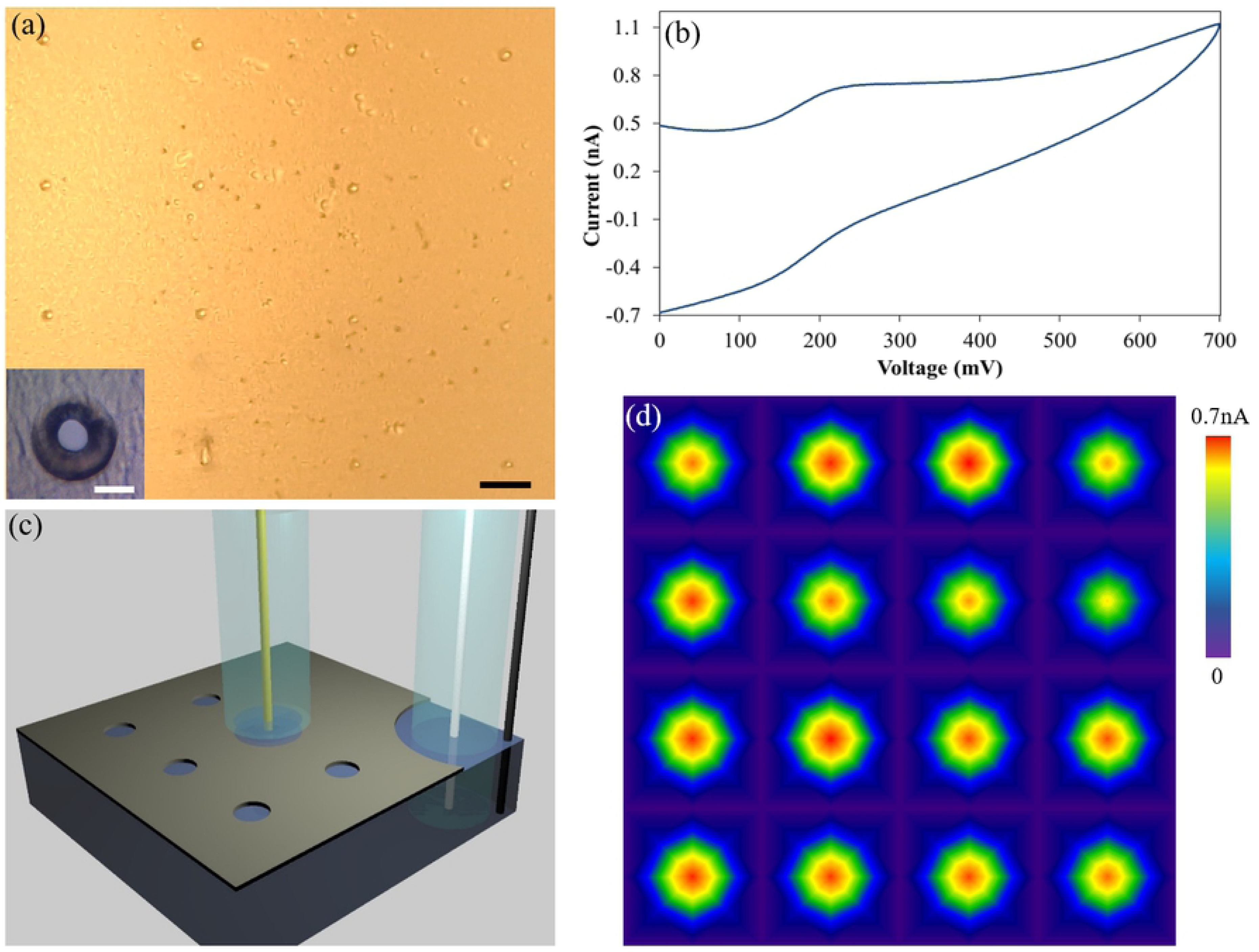
SECM setup testing in mapping electrochemical signals. (a) An image of the 4 by 4 array of pores on the parafilm membrane. Scale bar represents 1 mm. The average pore diameter was 80 micron as shown in the inset, where the scale bar depicts 100 micron. (b) A sample K_3_Fe(CN)_6_ cyclic voltammetry scan data from one of the pores in the array. (c) A computer rendered image of the experimental setup showing (not to scale) the working electrode over a pore and five other pores in the array. Reference and counter electrodes were immersed into the K_3_Fe(CN)_6_ solution as shown on the right corner. (d) A map of the cathodic current value at 200 mV obtained from the scan of 81 different points covering 1.2 cm by 1.2 cm area around the array of 16 pores.

In order to obtain the 4 by 4 array of pores, the motorized stage was utilized to move the etched tungsten wire with a pitch distance of 3 mm between pores, which was selected to match the diameter of the working microelectrode glass insulation tip diameter such that the working electrode doesn’t record electrochemical signal in between consecutive pores. The working electrode was a 12.5 micron diameter commercial gold microelectrode (CH Instruments, CHI105) in 3mm diameter glass insulation. Ag/AgCl reference electrode (CH Instruments, CHI111) and a 0.5 mm diameter platinum wire (Kurt Lesker) counter electrode were immersed into the K_3_Fe(CN)_6_ solution through a wider opening in the parafilm membrane further away from the 80 micron diameter pore array as depicted in Fig 3c. The motorized stage was used to move the working electrode with the GRBL software in scan steps of 1.5 mm to collect data from 81 different points in a 1.2cm x1.2cm area to cover all the pores in the array. In each step, the DSTAT potentiostat was used to record cyclic voltammograms of K_3_Fe(CN)_6_ solution (Fig 3b), which is in contact with the working electrode only through solution leakage from a pore as shown in the diagram in Fig 3c. The magnitude of the cathodic current at 200 mV was extracted from 81 data points, which was used in preparing the spatial electrochemical signal map around the pores as shown in Fig 3d. The cyclic voltammetry measurements yielded noise level signals in between pores as the trapped K_3_Fe(CN)_6_ solution under the working electrode didn’t contact the solution below the parafilm membrane at these locations. A variation in the signal amplitude was observed from pore to pore most likely due to non-uniformity of pore diameters. This result demonstrated the capability of the motorized stage in locating the pores precisely, which is necessary for proper functioning of the SECM.

## Biosensing test

In future experiments, we plan to use the SECM setup to scan an individual live cell surface with aptamer-based nanoscale electrodes for detecting target biomarkers released from the cell surface. The custom built SECM will enable the formation of submicron resolution spatial maps of the target biomarkers released from an individual cell.

As the custom built SECM will be used in future aptamer based biosensing experiments, the setup was also tested in DNA hybridization detection experiments. Voltammetry is commonly used in DNA hybridization experiments for the detection of target biomolecules [19,20]. Annealed tungsten wires with 250 micron diameter were used as working electrodes, which were tapered by electrochemical etching. The tapered tip tungsten electrodes were subsequently electroplated with gold by immersing in 20 mM Gold Chloride (HAuCl_4_) solution and by applying a 10V DC potential for 1 minute.

The gold coated electrode was rinsed with DI water and immersed in thiol functionalized 100 mM 20-nucleotide (20-nt) long ssDNA with poly Thymine (polyT) sequence for one hour. The polyT ssDNA was immobilized on the electrode through thiol-gold chemistry. The electrode was subsequently washed with PBS to eliminate unbound DNAs by immersing in 0.1 M PBS solution. Chronoamperometry measurements were performed with the DSTAT potentiostat by applying a 25mV vs Ag/AgCl constant potential and by measuring the current between the polyT ssDNA coated tungsten/gold electrode and a platinum reference electrode. A 30 μl aliquot of 100 mM 20-nt long ssDNA with poly Adenine (polyA) was added to the center of the container (∼1cm away from the working electrode) after 5 minutes of the experiment start time, where the current stabilized at ∼1.8 microamps following an initial characteristic RC drop as depicted by the orange line in Fig 4. In this experiment, the goal was the detection of polyA via monitoring the hybridization of polyA and polyT via Watson-Crick base-pairing. 20 minutes after the initiation of the experiment, the current started increasing steadily about three-fold compared to its starting value (from ∼4nA to ∼1.5 μA) in 10 minutes.

**Figure 4.**
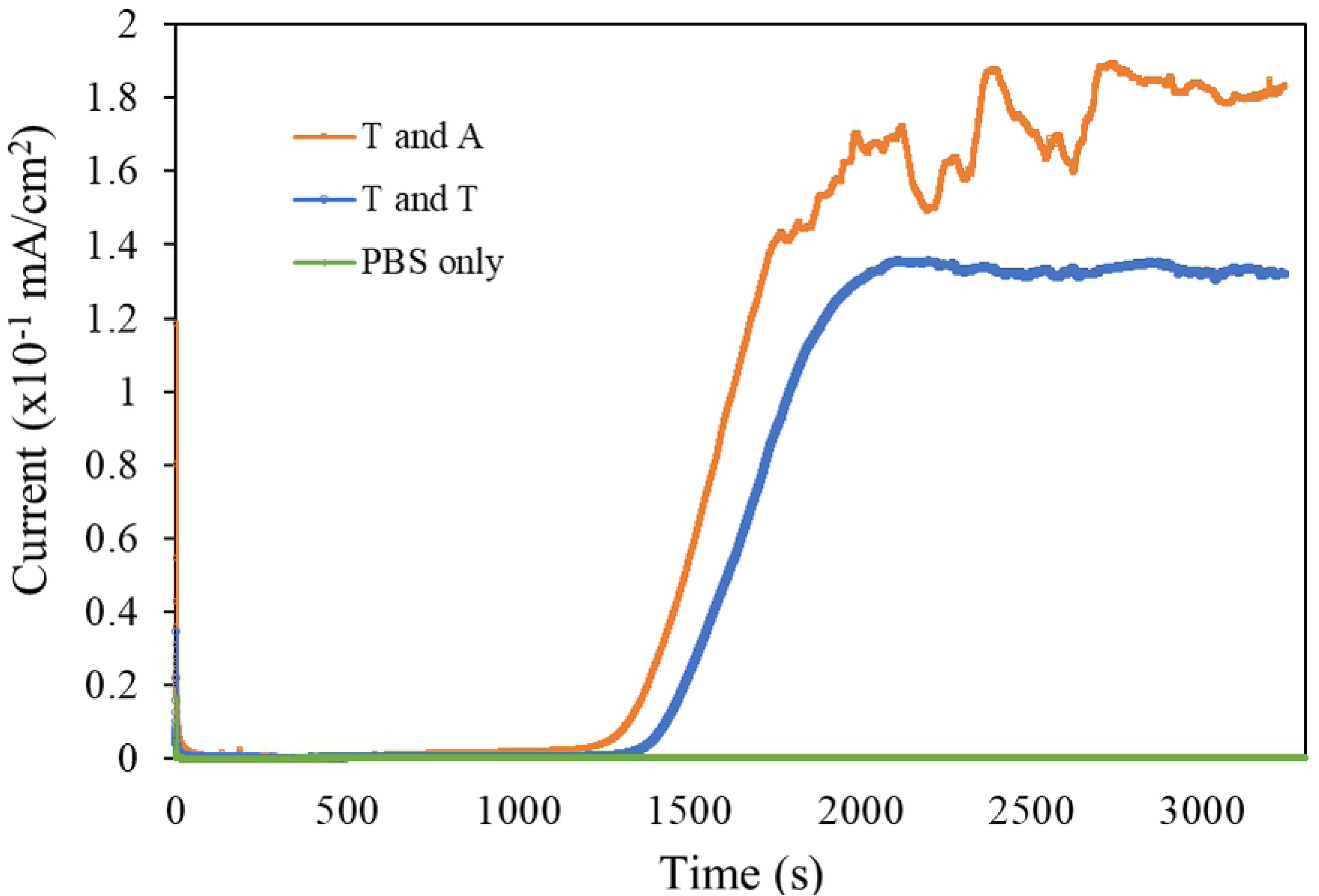
SECM setup testing in nucleic acid hybridization experiments. DNA hybridization measurements with chronoamperometry.

As a negative control, the experiment was repeated with a fresh polyT anchored electrode, where the same volume of 100 mM non-thiol functionalized 20-nt polyT was added to the fresh PBS solution. In this experiment, the current also increased steadily after about 22 minutes to similar elevated levels (blue line in Fig 4).

## Discussion

The steady current increase in both experiments was attributed to nonspecific adsorption of polyT and polyA onto the remaining available sites on the gold electrode surface. However, after the initial steady current increase in polyA detection experiment (orange line), current continued to increase at a reduced rate while fluctuating at various intervals for the rest of the experiment, a possible indication of a slower process (polyT and polyA hybridization) on the electrode surface. The negative control tested polyT (blue line) displayed a steady current after the first current increase step, which doesn’t hybridize with the thiolated polyT anchored electrode.

A second control experiment was performed by immersing a new gold coated tungsten electrode into the PBS solution. A steady base current was recorded for the whole duration of the experiment (green line), indicating that the current changes in the previous experiments were due to base pairing and nonspecific adsorption of target bases to the electrode surface.

Though it is not tested in the preliminary studies, the nonspecific adsorption observed with both polyT and polyA can be eliminated by backfilling the gold electrode surface with small thiol molecules such as 6-mercapto-1-hexanol (6-MCH). After immobilization of DNA on gold electrode through gold-thiol chemistry, addition of 6-MCH will inhibit adsorption of free ssDNAs to the electrode surface during the detection experiments. Because the 6-MCH is smaller than both the ssDNA and its 3’ or 5’ thiol linker, its influence on DNA: DNA hybridization is unlikely to happen. The surface passivation by 6-MCH will eventually eliminate the first-step current increase, and only the second-step current increase, which is a target specific detection event between polyT and polyA, will be measured in the future experiments. Nevertheless, the overall data demonstrated that the SECM setup has a strong potential for detection of biological materials; i.e., ssDNA, RNA, proteins and cancer biomarkers; though nucleic acid or aptamer binding.

## Conclusions

This work demonstrates the feasibility of building essential parts of a high performance SECM setup for less than $250 with DIY approach. The custom built SECM setup was tested by forming an electrochemical signal map from an array of pores in a membrane and by performing biosensing experiments. This SECM setup is not only a cost-efficient instrument but also its development is a hands-on training project for students on electronics, mechanics and electrochemistry. The modular nature of the setup also enables the utilization of individual components such as the potentiostat and the motorized stage for other experiments as needed. This custom-built SECM setup will be utilized in obtaining high-resolution electrochemical signal maps from individual cells with the use nanoscale electrodes in the future.

## Acknowledgments

We are thankful to Dr. Michael Dryden for providing technical support in the development and troubleshooting of DSTAT potentiostats. Research was sponsored by the Army Research Laboratory and was accomplished under Cooperative Agreement Number W911NF-12-2-0022. The views and conclusions contained in this document are those of the authors and should not be interpreted as representing the official policies, either expressed or implied, of the Army Research Laboratory or the U.S. Government. The U.S. Government is authorized to reproduce and distribute reprints for Government purposes notwithstanding any copyright notation herein. Research reported in this publication was partially supported by the National Institute of General Medical Sciences of the National Institutes of Health under Award Number UL1GM118973. The content is solely the responsibility of the authors and does not necessarily represent the official views of the National Institutes of Health.

## Supporting information

**S1_Supporting information. Additional hardware and software information.** Information on modifications to original DSTAT 3D printable box design, motorized stage parts and cost list, explanation of motorized stage 3D printable parts and instructions for GRBL software control of motorized stage are provided in this document.

**S1_Video. Demonstration of motorized stage submicron step capability.** The left part of the video shows a tapered tungsten wire tip on a microscope calibration slide. The tungsten wire is attached to the motorized stage. The distance between two lines is 10 microns. As the video starts, the user clicks the -Y button 20 times, which moves the tungsten wire tip for 10 microns in the -X direction on the screen. The alignment between the screen and the motion buttons can be corrected by rotating the microscope camera or by changing the orientation of the motorized stage relative to the microscope. The 0.045 mm step size on the GRBL screen (right part of the video) corresponds to 500 nm step motion by the motor, showing the submicron step size capability of the motorized stage.

**S2_Video. Automated pattern tracking of the motorized stage.** The tapered tungsten tip moves around the microscope calibration slide in a smiley face pattern according to a design provided by the user as a gcode file.

**S1_File. 3D printable part design files**. 3D printable stl files are provided including modified DSTAT box, motorized stage controller box, gears, and stepper motor brackets.

